# MerTK mediates the immunologically silent uptake of alpha-synuclein fibrils by human microglia

**DOI:** 10.1101/2023.03.04.531108

**Authors:** Marie-France Dorion, Konstantin Senkevich, Moein Yaqubi, Nicholas W. Kieran, Carol X.-Q. Chen, Adam MacDonald, Wen Luo, Amber Wallis, Irina Shlaifer, Jeffery A. Hall, Roy W. R. Dudley, Ian A. Glass, Birth Defects Research Laboratory, Jo Anne Stratton, Edward Fon, Tim Bartels, Jack P. Antel, Ziv Gan-or, Thomas M. Durcan, Luke M. Healy

## Abstract

MerTK is a receptor tyrosine kinase that mediates the immunologically silent phagocytic uptake of diverse types of cellular debris. Highly expressed on the surface of microglial cell, MerTK is of importance in brain development, homeostasis, plasticity, and disease. Yet, involvement of this receptor in the clearance of protein aggregates that accumulate with aging and in neurodegenerative diseases has yet to be defined. The current study explored the function of MerTK in the microglial uptake of alpha-synuclein fibrils which play a causative role in the pathobiology of synucleinopathies.

Using human primary and induced pluripotent stem cell-derived microglia, the MerTK- dependence of alpha-synuclein fibril internalization was investigated *in vitro*. Relevance of this pathway to synucleinopathies was assessed by analyzing MerTK expression in patient-derived cells and tissues.

Pharmacological inhibition of MerTK and siRNA-mediated *MERTK* knockdown both caused a decreased rate of alpha-synuclein fibril internalization by human microglia. Consistent with the immunologically silent nature of MerTK-mediated phagocytosis, alpha-synuclein fibril internalization did not induce secretion of pro-inflammatory cytokines from microglia. In addition, burden analysis in two independent patient cohorts revealed a significant association between rare functionally deleterious *MERTK* variants and Parkinson’s disease in one of the cohorts (*p* = 0.002). Accordingly, *MERTK* expression was significantly upregulated in nigral microglia from Parkinson’s disease/Lewy body dementia patients compared to those from non-neurological control donors in a single-nuclei RNA-sequencing dataset (*p* = 5.08*×*10^-21^), and MerTK protein expression positively correlated with alpha-synuclein level in human cortex lysates (*p* = 0.0029).

Taken together, our findings define a novel role for MerTK in mediating the uptake of alpha-synuclein aggregates by human microglia, with possible involvement in limiting alpha-synuclein spread in synucleinopathies such as Parkinson’s disease.

## Introduction

Synucleinopathies are neurodegenerative disorders characterized by the intracellular accumulation of aggregated alpha-synuclein (*α*-syn) protein. Broadly, they are classified into multiple system atrophy and Lewy body diseases that include Parkinson’s disease and Lewy body dementia. The neuronal accumulation of *α*-syn in the form of Lewy bodies and its role in neurodegenerative processes have been extensively studied. Duplication or triplication of the genetic locus encoding *α*-syn (*SNCA*) have been identified to cause autosomal dominant Parkinson’s disease,^1^ implying that accumulation of excess *α*-syn is sufficient to trigger the disease process. Current evidence supports the notion that the initiation of *α*-syn aggregation, either in the central nervous system or in peripheral tissues, can then propagate in a prion-like manner into the brain, where it induces neuronal dysfunction and degeneration.^2–7^ Accordingly, injection of recombinant *α*-syn preformed fibrils (PFFs) into mice leads to the development of Lewy body-like pathologies, neuronal loss and behavioral deficits.^8^ *In vitro,* seeding of neurons with *α*-syn PFFs also results in the formation of Lewy body-like inclusions and dysregulation of cellular processes.^9, 10^

Strategies to prevent cell-to-cell spread of *α*-syn have been proposed as potential therapies to prevent neurodegeneration in synucleinopathies.^11^ Mechanisms involved in *α*- syn endocytosis are being actively explored, with differential internalization mechanisms observed across brain cell types. For instance, heparan sulfate proteoglycans have been identified as mediators of *α*-syn PFF internalization by neurons but not microglia.^12^ Microglia as professional phagocytes residing in the brain can internalize and degrade *α*- syn fibrils with higher rate than other cells,^13–15^ making them a particularly promising cell type that can be targeted in the development of novel therapeutics to promote *α*-syn clearance. The majority of microglia studies investigating *α*-syn internalization processes to date have been carried out using murine microglia or the murine microglia-like cell line BV2, and have contributed to the identification of a variety of cell surface proteins that interact with human *α*-syn, including toll-like receptors (TLRs) and cluster of differentiation 36.^16–20^ Since human and murine microglia are inherently different in their genome, transcriptome and aspects of their function,^21–24^ it is unclear whether the same receptors and mechanisms participate in the uptake of *α*-syn by human microglia.

In a previous study comparing the impact of culture media formulation on microglia phagocytic activities, we observed a correlation between *α*-syn fibril and myelin debris uptake activities^25^ suggestive of a common mechanism of internalization for these substrates. Both *α*-syn fibril and myelin debris uptake activities also correlated with the expression level of Mer tyrosine kinase (MerTK), a microglial surface receptor kinase with known roles in myelin phagocytosis.^26^

<u>T</u>YRO3, <u>A</u>XL and <u>M</u>erTK constitute a family of homologous receptor tyrosine kinases called “TAM” with pleiotropic functions. AXL and MerTK are particularly known for their role in the phagocytosis of apoptotic cells,^27^ myelin debris,^26^ synapses,^28^ and photoreceptor outer segments.^29, 30^ Tethering of phagocytic targets to TAM receptors occurs through the binding of secreted bridging molecules – growth arrest-specific 6 (GAS6) and protein S1 (PROS1) – with high affinity for phosphatidylserine moieties exposed on the membranes of dead cells.^31^ Binding of GAS6/PROS1 to TAM receptors triggers activation of the cytoplasmic kinase domain, autophosphorylation and downstream signaling that induces cytoskeletal remodeling required for phagocytosis.^31^ This process is immunologically silent, and suppresses inflammatory activation of myeloid cells during homeostatic phagocytosis.^31, 32^

High expression of MerTK, as well as its activating ligands GAS6 and PROS1, is a characteristic feature of microglia that distinguishes them from other myeloid populations.^33^ The present study aimed to investigate the role of MerTK in *α*-syn fibril uptake by human primary (hMGL) and induced pluripotent stem cell-derived (iMGL) microglia. The relevance of this process in synucleinopathies was evaluated through rare variant burden analysis of *MERTK* in Parkinson’s disease and assessment of MerTK expression in patient-derived brain materials. Taken together, we show that *α*-syn fibril uptake by human microglia is largely MerTK-dependent. Increased MerTK expression accompanying *α*-syn accumulation in the human brain and genetic associations between rare *MERTK* variants and Parkinson’s disease suggest MerTK as a potential therapeutic target to enhance *α*-syn fibril clearance by microglia in synucleinopathies.

## Materials and Methods

### Ethical approval

Use of all human materials was approved by the McGill University Health Centre Research Ethics Board, under project# 1989-178 for human brain tissues and 2019-5374 for induced pluripotent stem cells (iPSCs).

### hMGL isolation and culture

Human brain tissues from 2- to 71-year-old female and male epilepsy patients were obtained from the Montreal Neurological Institute, Montreal, Canada (adult donors) and the Montreal Children’s Hospital, Montreal, Canada (pediatric donors), with written consent and under local ethic boards’ approval. Tissues were from sites distant from suspected primary epileptic foci. Second trimester fetal brain tissues were obtained from Centre Hospitalier Universitaire Sainte-Justine, Montreal, Canada or from the Birth Defects Research Laboratory, University of Washington, Seattle, USA with maternal written consent and under local ethic boards’ approval. Isolation of glial cells was carried out as previously described^34^ through mechanical and chemical digestion, followed by Percoll^®^ (Sigma-Aldrich) gradient centrifugation (postnatal tissue) or not (fetal tissue). hMGL were further purified by taking advantage of the differential adhesive properties of the glial cells. Cells of postnatal sources were maintained in microglia growth medium composed of Dulbecco’s Modified Eagle Medium (DMEM)/F12 (Thermo Fisher Scientific), 1% GlutaMAX^TM^ (Thermo Fisher Scientific), 1% non-essential amino acids (Thermo Fisher Scientific), 1% penicillin/streptomycin (P/S; Thermo Fisher Scientific), 2X insulin-transferrin-selenium (Thermo Fisher Scientific), 2X B27 (Thermo Fisher Scientific), 0.5X N2 (Thermo Fisher Scientific), 400 μM monothioglycerol (Sigma Aldrich) and 5% fetal bovine serum (FBS; Wisent Bioproducts), which was previously shown to promote high phagocytic activity and MerTK expression.^35^ Fetal hMGL were cultured in DMEM (Sigma-Aldrich) supplemented with 1% GlutaMAX^TM^, 1% P/S and 5% FBS. When indicated, hMGL were ‘M1’-polarized by treating them with 20 ng/mL interferon gamma (IFN*γ*; Peprotech) and 100 ng/mL Pam3CSK4 (Invivogen) for 48 hours or left unpolarized (‘M0’). Cells were maintained at 37 °C under a 5% CO2 atmosphere.

### Generation of iMGL

A number of iPSC lines were used in our study to ensure reproducibility of our findings across a range of different genetic backgrounds: DYR0100 (American Type Cell Collection), GM25256 (Coriell Institute) and 3450 (generated in-house).^36^ Differentiation of iPSC into iMGL was carried out following a previously established protocol.^37^ Briefly, hematopoietic progenitor cells were generated from iPSC using STEMdiff Hematopoietic kit (STEMCELL Technologies) and cultured in microglia growth medium supplemented with 100 ng/mL interleukin-34, 50 ng/mL tumor growth factor-beta and 25 ng/mL macrophage colony-stimulating factor (Peprotech) for 25 days, following which 100 ng/mL cluster of differentiation 200 (Abcam) and C-X3-C motif chemokine ligand 1 (Peprotech) were also added to the culture. Cells were maintained at 37 °C under a 5% CO2 atmosphere throughout the protocol.

### RNA-sequencing (RNAseq)

RNAseq data were obtained from an earlier study.^25^ Quality control of the RNA samples, the library preparation and RNA-sequencing were performed by Genome Quebec, Montreal, Canada. The library was generated using a NEBNext® Single Cell/Low Input RNA Library Prep Kit (New England Biolabs). RNAseq was performed using an Illumina NovaSeq 6000. Canadian Center for Computational Genomics’ pipeline GenPipes^38^ was used to align the raw files and quantify the read counts.

### Western blotting

Cells were lysed on ice in a lysis buffer composed of 150 mM NaCl, 50 mM Tris-HCl pH 7.4, 1% Nonidet P-40, 0.1% sodium dodecyl sulfate and 5 mM ethylenediaminetetraacetic acid with protease and phosphatase inhibitors (Thermo Fisher Scientific). Cell lysates were centrifuged at 500 g for 30 minutes at 4°C to remove cellular debris. Proteins (25 μg/lane) were separated on SDS-polyacrylamide gels and transferred to polyvinylidene difluoride membranes (Bio-Rad Laboratories). For the detection of *α*- syn, membranes were fixed in 4% paraformaldehyde solution as previously described.^39^ Membranes were immunoblotted for AXL (AF154 R&D Systems at 1:500), MerTK (ab52968, Abcam at 1:500), *α*-syn (ab138501, Abcam 1:5000) and glyceraldehyde 3- phosphate dehydrogenase (GAPDH; G8795, Sigma Aldrich at 1:5000) overnight at 4°C, and then with horse radish peroxidase-linked secondary antibodies (1:10000; Jackson Laboratory) for one hour. Bands were detected by enhanced chemiluminescence with Clarity Max ECL substrates (Bio-Rad Laboratories) using a ChemiDoc Imaging System (Bio-Rad Laboratories). Image analysis was performed using ImageLab 6.0.1 software (Bio-Rad Laboratories).

### Preparation of *α*-syn PFFs

PFFs of *α*-syn were generated as previously described^40, 41^ with slight modifications. Briefly, glutathione-S-transferase (GST) -tagged human recombinant *α*-syn were expressed and isolated from *Escherichia coli*. The GST tag was removed and purified *α*-syn monomers were subjected to endotoxin removal using Pierce^TM^ High-Capacity Endotoxin Removal Resin (Thermo Fisher Scientific) to achieve endotoxin levels < 0.3 EU/mg. *α*-syn monomers were aggregated on a shaker for five days with constant shaking at 1000 rpm, at 37°C. Resulting PFFs were subjected to 40-80 cycles of sonication (30 seconds ON/30 seconds OFF) using a Zetasizer Nano (Malvern Panalytical) to achieve *∼*50-100 nm sizes. Sizes and morphology of PFFs were monitored by electron microscopy and dynamic light scattering. Thioflavin T assay was used to confirm presence of beta-sheet structure. All data were validated using *α*-syn PFFs from at least two different batches.

### Uptake assay

Human *α*-syn PFFs, myelin debris^26^ and immunoglobulin G-opsonized red blood cells^42^ (IgG-RBCs) were labelled with pHRodo Green^TM^ STP ester (Thermo Fisher Scientific) and were used at the following respective concentrations which were determined to be non-saturating: 1 μM, 15 μg/mL and 50 000 cells/mL respectively. Cells were incubated with the labelled substrates for two hours and then counterstained with Hoechst 33342 (5 μg/mL). Total green fluorescence intensity per cell was quantified using a CellInsight CX5 High Content Screening Platform (Thermo Fisher Scientific). All conditions were assessed in triplicate. Unchallenged cells were used to measure background/autofluorescence. Internalization of Alexa Fluor 488-conjugated human *α*-syn PFFs (1 μM) and epidermal growth factor (EGF; 4 μg/mL; Thermo Fisher Scientific) were assessed similarly, except cells were washed with 4% trypan blue solution to quench extracellular fluorescence. When indicated, cells were pretreated for one hour with non-cytotoxic concentrations of UNC2025 (3 μM; Cayman Chemical) or cytochalasin D (1 μM; Sigma Aldrich).

### RNA interference

Cells were transfected with siGENOME RISC-Free Control (siCON) or ON-TARGETplus small interfering RNA (siRNA) pools against *MERTK* (siMERTK, at 20 nM) and/or *AXL* (siAXL, at 10 nM) (Horizon Discovery) using Lipofectamine^TM^ RNAiMAX (Thermo Fisher Scientific). The manufacturers’ protocols were followed.

When necessary, siCON was used to equilibrate the total amount of siRNA received by cells across experimental conditions.

### Flow cytometry

Cells were blocked with Human TrueStain FcX and TrueStain Monocyte Blocker (Biolegend) and stained with the following antibodies: anti-AXL clone #108724 and anti-MERTK clone #125518 (R&D Systems). Appropriate forward and side scatter profiles were used to exclude debris and doublets from the analysis. Dead cells were excluded based on LIVE/DEAD^TM^ Fixable Aqua (Thermo Fisher Scientific) staining. Readings were done on an Attune^TM^ Nxt Flow Cytometer and analyzed/visualized using FlowJo^TM^ software.

### Cell viability

Cell viability was assessed by propidium iodide (PI; Thermo Fisher Scientific) staining unless otherwise specified. Cells were stained in phosphate buffered saline containing 1 μg/mL PI and 5 μg/mL Hoechst 33342 (Thermo Fisher Scientific) and the average number of PI- live cells per condition was determined using a CellInsight CX5 High Content Screening Platform. All conditions were assessed in triplicate.

### Quantitative reverse transcription polymerase chain reaction (qRT-PCR)

A RNeasy mini kit (Qiagen) was used to extract RNA. Reverse transcription was performed using Moloney murine leukemia virus reverse transcriptase (Thermo Fisher Scientific) and real-time PCR was performed using TaqMan assays (Thermo Fisher Scientific) on a QuantStudio^TM^ 5 real-time PCR system (Thermo Fisher Scientific). The 2^- ΔCt^ method was used to analyze the data using *GAPDH* and tyrosine 3- monooxygenase/tryptophan 5-monooxygenase activation protein zeta (*YWHAZ*) as controls.

### Measurement of cytokine secretion

Concentrations of interleukin-1beta (IL-1*β*), interleukin-6 (IL-6), interleukin-10 (IL-10) and tumor necrosis factor (TNF) in cell supernatants were measured using the Human Inflammatory Cytokine Cytometric Bead Array Kit (BD Biosciences). Readings were done on an Attune^TM^ Nxt Flow Cytometer.

### Burden analysis of rare variants

Rare variant analysis was performed in the UK Biobank (UKBB) cohort, composed of 602 Parkinson’s disease patients and 15,000 randomly sampled controls, and the Accelerating Medicines Partnership – Parkinson Disease (AMP-PD) initiative cohort, composed of 2,341 Parkinson’s disease patients and 3,486 controls. All Parkinson’s disease patients were diagnosed according to either the UK Brain Bank criteria^43^ or the Movement Disorders Society criteria.^44^ Only individuals with European ancestry were included. For the UKBB cohort, whole-exome sequencing data was used. Quality control was performed both at the individual and variant levels using Genome Analysis Toolkit (GATK, v3.8) and plink, with a minimum depth of coverage 10 and GQ = 20 as described previously.^45^ For the AMP-PD cohort, we used all available whole-genome sequencing data for which the quality control processes at the individual and variant levels have been previously described (https://amp-pd.org/whole-genome-data). Associations between variants and Parkinson’s disease was tested using optimized sequence Kernel association test (SKAT-O).^46^ Variants were categorized into a) rare (minor allele frequency <1%) nonsynonymous variants, b) functional variants (nonsynonymous, stop/frameshift and splicing) and c) variants with a combined annotation dependent depletion (CADD) score of ≥ 20, representing the top 1% of potentially deleterious variants. A meta-analysis of the two cohorts was run using the metaSKAT package.^47^

### Analysis of single-nuclei RNA-sequencing (snRNAseq) data

snRNAseq data of human substantia nigra were from Kamath *et al.*, 2022,^48^ and were accessed through GEO:178265. Dataset included eight non-neurological control donors, seven Parkinson’s disease patients and three Lewy body dementia patients. All patients were pathologically confirmed to present moderate to prominent loss of pigmentation in the substantia nigra.^48^ Microglia population was subtracted from the main object using metadata information provided by the authors at the Broad Institute Single Cell Portal https://singlecell.broadinstitute.org/single_cell/study/SCP1768/. The subtracted microglia population was subjected to the standard Seurat pipeline for dimensionality reduction, gene expression normalization, clustering, and differential expression analysis. Twenty clusters expressing microglia canonical markers were identified (Supplementary Fig. 1), among which clusters 8, 12, 16 and 17 showed enrichment in stress response genes typically associated with tissue dissociation procedures (Supplementary Table 1). Those clusters, along with cluster 11 with macrophage marker expression (Supplementary Table 1), were removed from further analysis. Differentially expressed genes (DEGs) between Parkinson’s disease/Lewy body dementia and control microglia were identified using *ç*log2(fold change)*÷* > 0.25 and adjusted p-value < 0.05 as cutoffs.

### Preparation of human cortex lysates

Human cortex tissues were from the Mayo Clinic Brain Bank, Jacksonville, USA. Tissues were derived from five Lewy body dementia patients and five control donors devoid of Lewy body disease. Tissues were dounce-homogenized in Tris-buffered saline with protease and phosphatase inhibitors. Samples were further diluted in lysis buffer with 5% sodium dodecyl sulfate, protease and phosphatase inhibitors for protein extraction as described above. Clinicopathological information of the donors is presented in Supplementary Table 2.

### Statistical analyses

Statistical analyses were performed using GraphPad Prism 9.0 software. A t-test was used to compare the mean of two groups of data. A one-way analysis of variance (ANOVA) was used to compare the mean of three or more groups of data. When the assumptions of a t-test or a one-way ANOVA were not met, a Mann-Whitney test or a Kruskal-Wallis test was used instead, respectively. P-values (‘*p*’) were adjusted using appropriate post hoc tests following one-way ANOVA/Kruskal-Wallis tests. Mean and standard error of the mean (SEM) of biological replicates (‘*n*’) are plotted in all graphs unless otherwise indicated. A *p* < 0.05 was considered statistically significant. hMGL obtained from independent donors and iMGL generated at different points in time were considered biological replicates.

### Data availability

RNAseq data of hMGL and iMGL are available in Dorion *et al.*, 2023.^25^ snRNAseq data from Kamath *et al.*, 2022^48^ are available through Gene Expression Omnibus (<u>GSE178265</u>).

## Results

### hMGL and iMGL express MerTK and internalize *α*-syn PFFs

We first aimed to assess the MerTK-dependence of *α*-syn PFF uptake by human microglia. Given the limited access to hMGL, the potential of iMGL as an alternative model to investigate the role of MerTK in *α*-syn PFF uptake was evaluated. RNAseq and Western blotting of TAM receptor expression revealed that both AXL and MerTK are expressed in hMGL and iMGL (Figure 1A-D). Interestingly, AXL expression was significantly higher in iMGL compared to hMGL, whereas MerTK expression was similar between both cell types (Figure 1A-D). As expected, *TYRO3* expression was low in both models (Figure 1A-B). Given these results, subsequent experiments were carried out in iMGL, with validation of key findings in hMGL.

**Figure 1.**
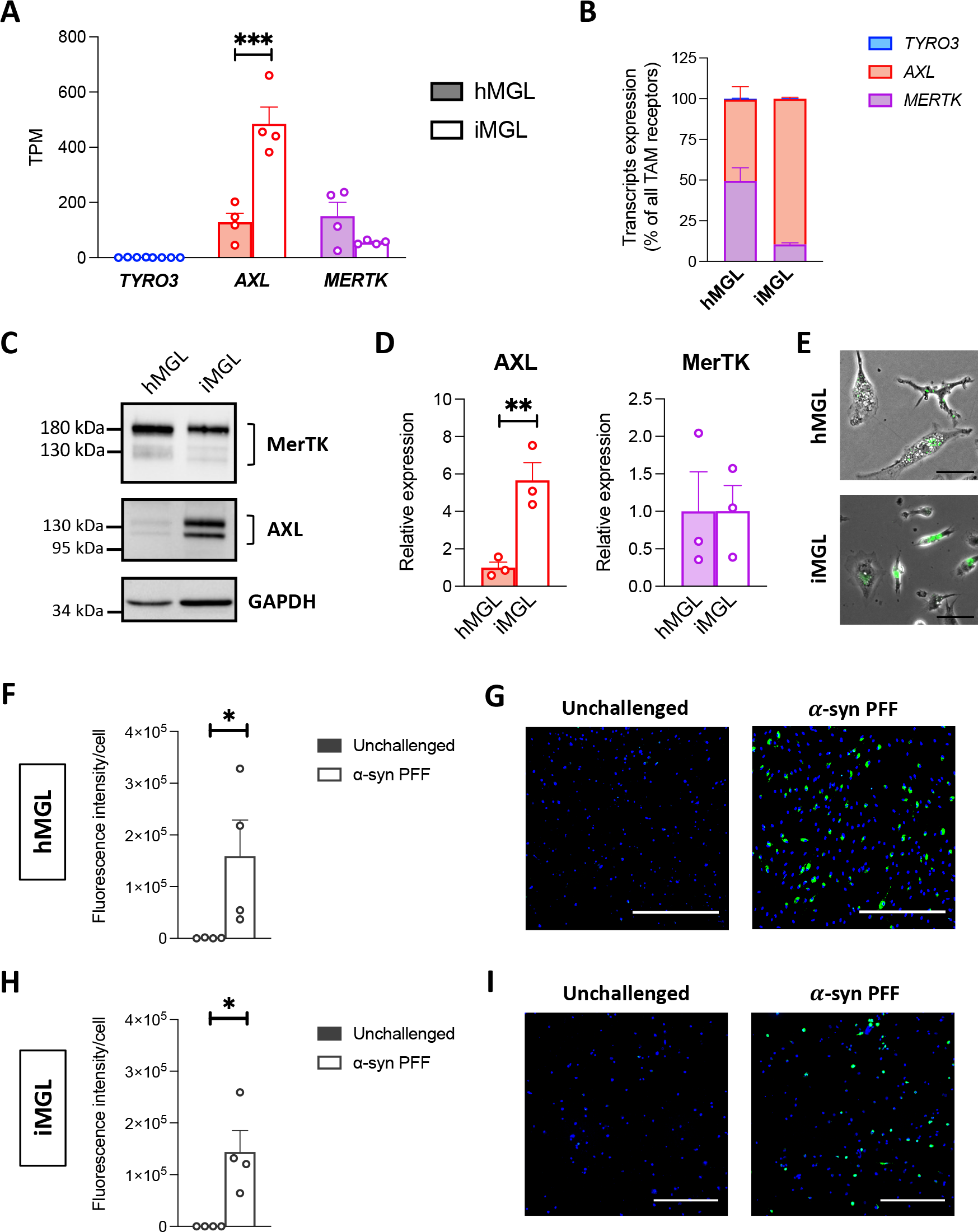
TAM receptors expression and *α*-syn PFF uptake by human microglia. (A- B) RNAseq assessment of *TYRO3*, *AXL* and *MERTK* expression in hMGL and iMGL. A one-way ANOVA with Sidak’s post hoc test was performed for pairwise comparisons between hMGL and iMGL in (A). TPM = transcripts per million. Mean +/- SEM of *n* = 4, \*\*\**p* < 0.001. (C) Western blot of MerTK, AXL and GAPDH. (D) Quantification of AXL and MerTK expression in iMGL relative to hMGL. GAPDH was used as a loading control. T-tests were performed. Mean +/- SEM of *n* = 3, \*\**p* < 0.01. (E-I) hMGL and iMGL were challenged with pHrodo^TM^ Green-labelled *α*-syn PFFs for two hours. (E) Merged phase contrast and green fluorescence images. Scale bar = 50 μm. (F) Quantification of green fluorescence intensity per cell in hMGL culture. A Mann-Whitney test was performed. Mean +/- SEM of *n* = 4, \**p* < 0.05. (G) Representative fluorescence images of hMGL counterstained with Hoechst 33342 (blue). Scale bar = 500 μm. (H) Quantification of green fluorescence intensity per cell in iMGL culture. A Mann-Whitney test was performed. Mean +/- SEM of *n* = 4, \**p* < 0.05. (I) Representative fluorescence images of iMGL counterstained with Hoechst 33342 (blue). Scale bar = 300 μm.

pHrodo^TM^ dyes are fluorogenic dyes that fluoresce when present in an acidic milieu such as phagosomes and lysosomes; and have been previously demonstrated to be useful tools for quantifying the rate of phagocytosis.^26, 49^ After a two-hour incubation of iMGL with pHrodo^TM^ Green-labelled *α*-syn PFFs, a dose-dependent increase in green fluorescence intensity per cell was quantified (Supplementary Fig. 2). A concentration of 1 μM was non-saturating (Supplementary Fig. 2) and resulted in readily quantifiable fluorescence intensity in both hMGL (Figure 1E-G) and iMGL cultures (Figure 1E, H-I).

### Pharmacological inhibition of MerTK results in decreased *α*- syn PFF uptake by microglia

The selective MerTK inhibitor UNC2025^50^ was used to investigate the involvement of MerTK in *α*-syn fibril internalization by microglia. A one-hour pretreatment with UNC2025 reduced iMGL uptake of pHrodo^TM^ Green-labelled *α*-syn PFFs by *∼*79% (Figure 2A-C). UNC2025 also decreased pHrodo^TM^ Green-labelled myelin uptake as previously described,^26^ but had no effect on pHrodo^TM^ Green-labelled IgG-RBC uptake, a process mediated by Fc gamma receptors and thus independent of MerTK^27, 51^ (Figure 2A- C). In contrast, the actin polymerization inhibitor cytochalasin D non-selectively inhibited the internalization of all three substrates (Figure 2A-C). UNC2025 also decreased hMGL uptake of pHrodo^TM^ Green-labelled *α*-syn PFFs by *∼*72% (Figure 2D-E), providing validation that MerTK kinase activity is essential for the internalization of *α*-syn fibrils by hMGL.

**Figure 2.**
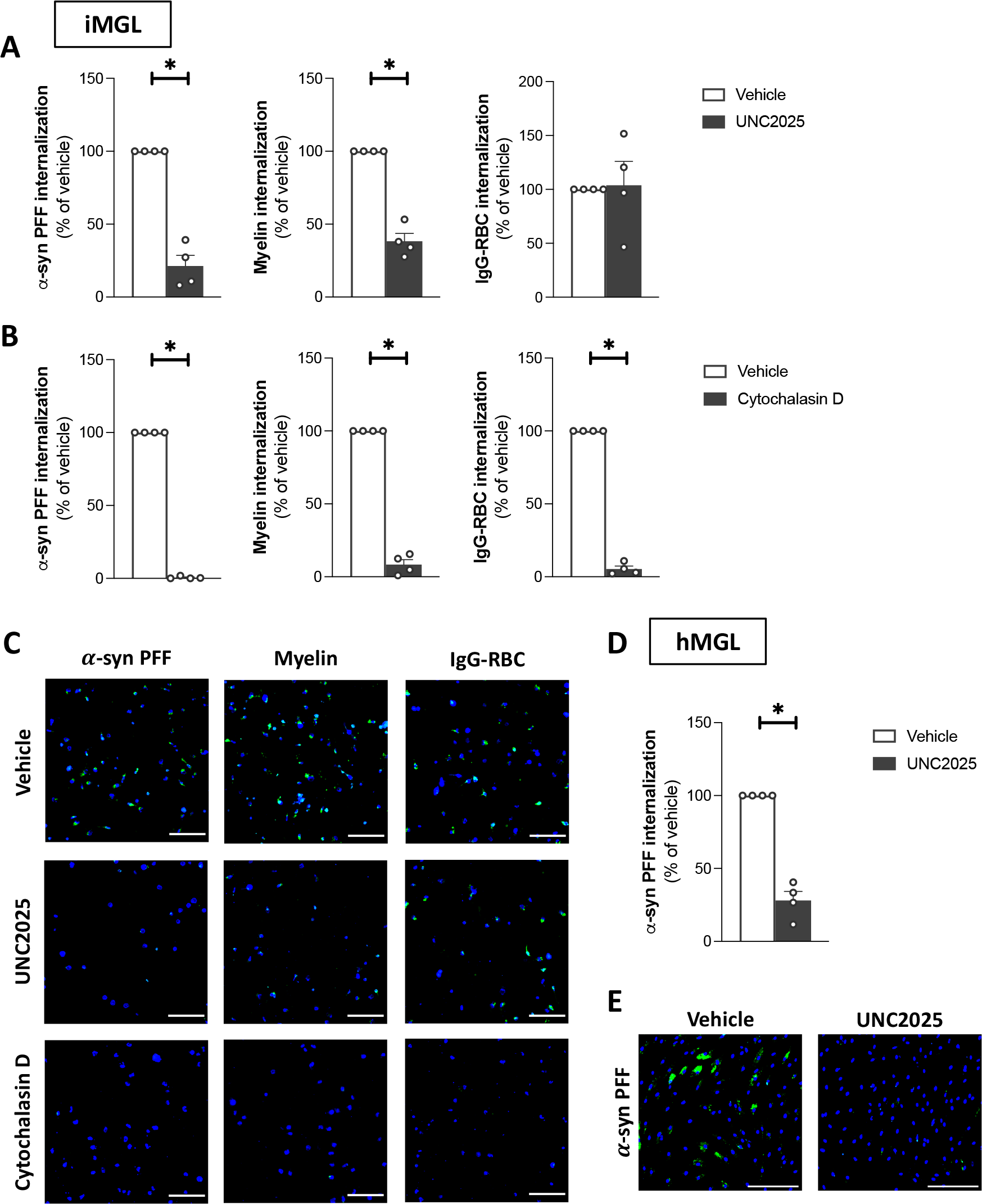
Effect of UNC2025 on microglial uptake of *α*-syn PFFs. Microglia were pretreated with vehicle, UNC2025 (3 *μM*) or cytochalasin D (1 *μM*) for one hour, then challenged with pHrodo^TM^ Green-labelled *α*-syn PFFs, myelin debris or IgG-RBCs for two hours. (A-B) Quantification of green fluorescence intensity per iMGL. Data were normalized to the vehicle-treated conditions. A Mann-Whitney test was performed. Mean +/- SEM of *n* = 4, \**p* < 0.05. (C) Representative images of iMGL counterstained with Hoechst 33342 (blue). Scale bar = 150 *μm*. (D) Quantification of green fluorescence intensity per hMGL. Data were normalized to the vehicle-treated conditions. A Mann- Whitney test was performed. Mean +/- SEM of *n* = 4, \**p* < 0.05. (E) Representative images of hMGL counterstained with Hoechst 33342 (blue). Scale bar = 200 *μm*.

To ensure that results were not influenced by the use of a pH-sensitive dye, additional tests were performed with Alexa Fluor 488-labelled *α*-syn PFFs. Similar to findings with pHrodo^TM^ Green-labelled *α*-syn PFFs, UNC2025 also decreased the internalization of Alexa Fluor 488-labelled *α*-syn PFFs by iMGL (by ∼79%; Supplementary Fig. 3A-B), confirming that MerTK appears to mediate the internalization of *α*-syn PFFs regardless of the conjugated dyes. UNC2025 did not affect the internalization of Alexa Fluor 488- labelled EGF which is known to be mediated by epidermal growth factor receptor^52^ (EGFR; Supplementary Fig. 3A-B). All subsequent uptake assays were performed using pHrodo^TM^ Green-labelled *α*-syn PFFs.

### *MERTK* knockdown results in decreased *α*-syn PFF uptake by microglia

siRNA-mediated knockdown experiments were next carried out to confirm that our earlier findings were a consequence of MerTK inhibition and not an off-target effect of UNC2025 on other tyrosine kinases. iMGL were transfected with siRNAs against *AXL* and/or *MERTK*, and replated for the assessment of *α*-syn PFF uptake. siAXL and siMERTK significantly decreased the respective surface expression of AXL and MerTK on live iMGL after 48 hours (Figure 3A). Knockdown of *AXL* had no marked impact on *α*-syn PFF internalization by iMGL, unlike *MERTK* knockdown which resulted in a *∼*71% decrease in internalization (Figure 3B-C). Knockdown of both *AXL* and *MERTK* together did not further affect *α*-syn PFF uptake compared to *MERTK* knockdown alone (Figure 3B-C). Similarly, in hMGL, *AXL* knockdown did not affect *α*-syn PFF uptake, whereas *MERTK* knockdown did (by *∼*56%; Figure 3D-F). Neither *AXL* nor *MERTK* knockdown affected hMGL viability (Figure 3G), ruling out the possibility that a viability effect is downregulating PFF internalization. Altogether, these findings imply that MerTK but not AXL is essential for *α*-syn fibril uptake by human microglia.

**Figure 3.**
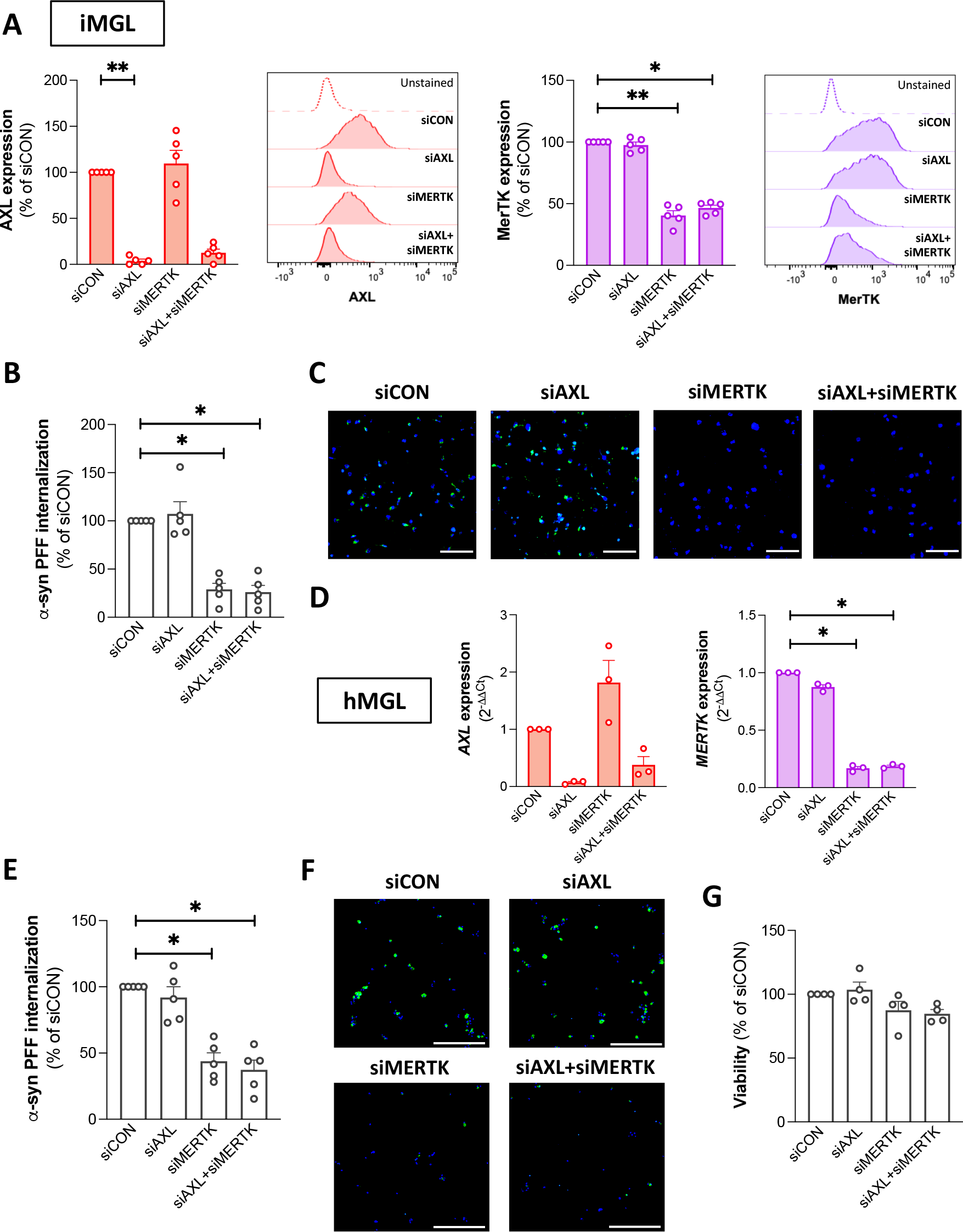
Effect of *AXL*/*MERTK* knockdown on microglial uptake of *α*-syn PFFs. Microglia were transfected with siCON, siAXL and/or siMERTK. (A) Cell surface expression of AXL and MerTK assessed by flow cytometry on live iMGL 48 hours following transfection. Data were normalized to the siCON-transfected conditions. Kruskal-Wallis tests were performed followed by Dunn’s post hoc test. Mean +/- SEM of *n* = 5, \**p* < 0.05, \*\**p* < 0.01. (B) Quantification of green fluorescence intensity per cell in iMGL culture challenged with pHrodo^TM^ Green-labelled *α*-syn PFFs for two hours. Data were normalized to the siCON-transfected conditions. A Kruskal-Wallis test was performed followed by Dunn’s post hoc test. Mean +/- SEM of *n* = 5, \**p* < 0.05. (C) Representative images of iMGL challenged with pHrodo^TM^ Green-labelled *α*-syn PFFs for two hours and counterstained with Hoechst 33342 (blue). Scale bar = 150 *μm*. (D) qRT- PCR assessment of *AXL* and *MERTK* expression in hMGL 48 hours following transfection. Kruskal-Wallis tests were performed followed by Dunn’s post hoc test. Mean +/- SEM of *n* = 3, \**p* < 0.05. (E) Quantification of green fluorescence intensity per cell in hMGL culture challenged with pHrodo^TM^ Green-labelled *α*-syn PFFs for two hours. Data were normalized to the siCON-transfected conditions. A Kruskal-Wallis test was performed followed by Dunn’s post hoc test. Mean +/- SEM of *n* = 5, \**p* < 0.05. (F) Representative fluorescence images of hMGL challenged with pHrodo^TM^ Green-labelled *α*-syn PFFs for two hours and counterstained with Hoechst 33342 (blue). Scale bar = 200 *μm*. (G) Viability of hMGL 48 hours following transfection. Kruskal-Wallis tests were performed followed by Dunn’s post hoc test. Mean +/- SEM of *n* = 3, \**p* < 0.05.

Previously, it has been shown that combined stimulation of microglia with IFN*γ* and a TLR agonist, the so-called “M1” polarization, results in a pro-inflammatory phenotype characterized by a decreased expression of MerTK and a lower internalization of myelin debris.^26^ Polarization of hMGL with IFN*γ* and the TLR2/1 agonist Pam3CSK4 was confirmed to increase the M1 markers human leukocyte antigen-DR (*HLA-DR*), *TNF* and C-X-C motif chemokine ligand 10 (*CXCL10*) expression, and to decrease *IL10* expression compared to the unpolarized M0 state (Supplementary Fig. 4A). Expression of genes encoding MerTK and its activating ligands GAS6 and PROS1 were decreased in M1 cells whereas AXL expression was unchanged (Supplementary Fig. 4A). This decreased expression of *MERTK*, *GAS6* and *PROS1* was associated with significantly lower uptake of *α*-syn PFFs (Supplementary Fig. 4B-C), further supporting the notion that MerTK alone, but not AXL mediates *α*-syn fibril uptake by human microglia.

### MerTK-mediated *α*-syn PFF internalization is immunologically silent

Phagocytic processes involving MerTK are known to be immunologically silent.^53^ With pharmacological and knockdown assays pointing towards MerTK being a central mediator of *α*-syn fibril uptake, the effects of *α*-syn PFFs on immune pathways was investigated next. The widely used inflammatory stimulant and TLR agonist Pam3CSK4 significantly increased the secretion of IL-6 and TNF by iMGL, confirming that these cells can be activated to secrete cytokines (Figure 4A). Consistent with the idea that MerTK-mediated phagocytosis is immunologically silent, a 24-hour treatment with *α*-syn PFFs did not result in increased iMGL secretion of IL-6, TNF, IL-10 or IL-1*β* (Figure 4A). Interestingly, a modest but statistically significant decrease in the secretion of those cytokines was observed following *α*-syn PFF treatment (Figure 4B), indicative of an immunomodulatory effect. The *α*-syn PFF-induced decrease in IL-6, TNF and IL-1*β* was prevented by a one-hour pretreatment of iMGL with UNC2025 or cytochalasin D (Figure 4B), suggesting that this immunomodulatory effect is dependent on MerTK-mediated *α*- syn PFF internalization. The decrease in IL-10 was not prevented by UNC2025 or cytochalasin D pretreatment (Figure 4B), suggesting that this effect of *α*-syn PFFs is independent from their internalization. Treatment of hMGL with *α*-syn PFFs also did not elicit an increase in IL-6, TNF, IL-10 or IL-1*β* secretion (Figure 4C). IL-1*β* secretion was significantly inhibited by *α*-syn PFFs in the hMGL (Figure 4C).

**Figure 4.**
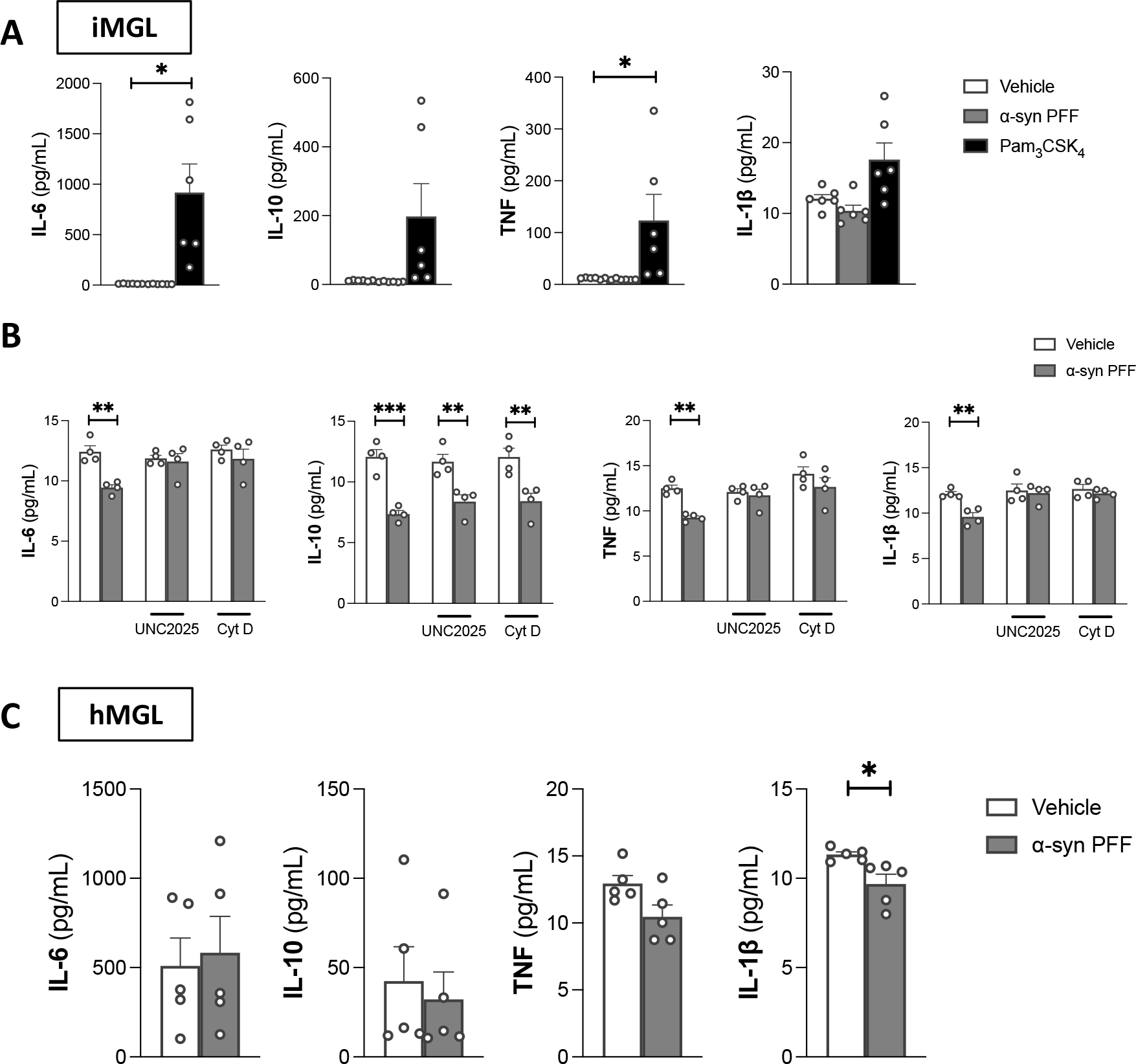
Inflammatory response of human microglia to *α*-syn PFFs. Microglia were treated for 24 hours with vehicle, *α*-syn PFFs (1 *μM*) or Pam3CSK4 (100 ng/mL). (A) Cytokine secretion from iMGL. Kruskal-Wallis tests were performed followed by Dunn’s post hoc test. Mean +/- SEM of *n* = 6, \**p* < 0.05. (B) Cytokine secretion from iMGL pretreated or not with UNC2025 (3 *μM*) or cytochalasin D (cyt D; 1 *μM*) for one hour prior to vehicle/*α*-syn PFF treatment. Kruskal-Wallis tests were performed followed by Dunn’s post hoc test. Mean +/- SEM of *n* = 4, \*\**p* < 0.01, \*\*\**p* < 0.001. (C) Cytokine secretion from hMGL. Kruskal-Wallis tests were performed followed by Dunn’s post hoc test. Mean +/- SEM of *n* = 6, \**p* < 0.05.

Expression of a wider range of cytokines was assessed in iMGL by qRT-PCR following three hours, one day, and seven days of *α*-syn exposure. In contrast to Pam3CSK4, neither the monomeric nor fibrillar forms of *α*-syn increased the transcriptional expression of cytokines (Supplementary Fig. 5). To summarize, *α*-syn did not induce an inflammatory response of human microglia similar to that triggered by TLR stimulation.

### Rare deleterious *MERTK* variants are associated with Parkinson’s disease

To further examine the potential role of MerTK-mediated phagocytosis in synucleinopathies, genetic associations between rare *MERTK* variants and Parkinson’s disease were investigated by burden analysis in two independent patient cohorts: UKBB and AMP-PD. A significant association between *MERTK* variants with CADD score ≥ 20 (representing the top 1% of potentially deleterious variants) and Parkinson’s disease was found in the UKBB cohort (*p* = 0.002, Table 1). However, this association was not observed in the AMP-PD cohort or following a meta-analysis of both cohorts (Table 1). No significant associations were found between *AXL* variants and Parkinson’s disease (Table 1). These results suggest that some rare, functionally deleterious *MERTK* variants may be associated with Parkinson’s disease, however these findings require additional replications.

**Table 1.** Burden analysis of rare variants in Parkinson’s disease

| Gene | All nonsynonymous, $p$ | | | All functional, $p$ | | | CADD score $\geq 20$, $p$ | | |
| --- | --- | --- | --- | --- | --- | --- | --- | --- | --- |
|  | UKBB | AMP-PD | Meta-analysis | UKBB | AMP-PD | Meta-analysis | UKBB | AMP-PD | Meta-analysis |
| *AXL* | 0.84 | 0.27 | 0.48 | 0.83 | 0.27 | 0.48 | 0.45 | 0.85 | 0.54 |
| *MERTK* | 0.2 | 0.92 | 0.81 | 0.21 | 1 | 0.78 | **0.002** | 0.67 | 0.28 |

### *MERTK* expression is upregulated in nigral microglia of Parkinson’s disease patients

Following the observation that rare genetic variants of *MERTK* are potentially associated with Parkinson’s disease, the possibility that the MerTK pathway is dysregulated in the Parkinson’s disease substantia nigra was investigated using a previously published postmortem snRNAseq dataset.^48^ DEG analysis of microglia from Parkinson’s disease (*n* = 7) or Lewy body dementia (*n* = 3) substantia nigra tissues compared to those of non-neurological controls (*n* = 8) revealed a decrease in expression of the homeostatic markers C-X3-C motif chemokine receptor 1 (*CX3CR1*) and transforming growth factor beta receptor 1 (*TGFBR1*), accompanied by an increase in the expression of specific neurodegeneration-associated genes that included glycoprotein NMB (*GPNMB*) and galectin-3 (*LGALS3*), in the diseased group (Figure 5A). Expression of *SNCA* and leucine-rich repeat kinase 2 (*LRRK2*) with known genetic association to Parkinson’s disease was also significantly elevated in microglia from Parkinson’s disease/Lewy body dementia patients (Figure 5A). Consistent with our *in vitro* data, genes encoding inflammatory cytokines did not figure among the DEGs (Supplementary Table 3). Most importantly, evaluation of genes encoding TAM receptors and their activating ligands revealed *MERTK* expression to be significantly higher in microglia from diseased patients compared to controls (*p* = 5.08*×*10^-21^; Figure 5B).

**Figure 5.**
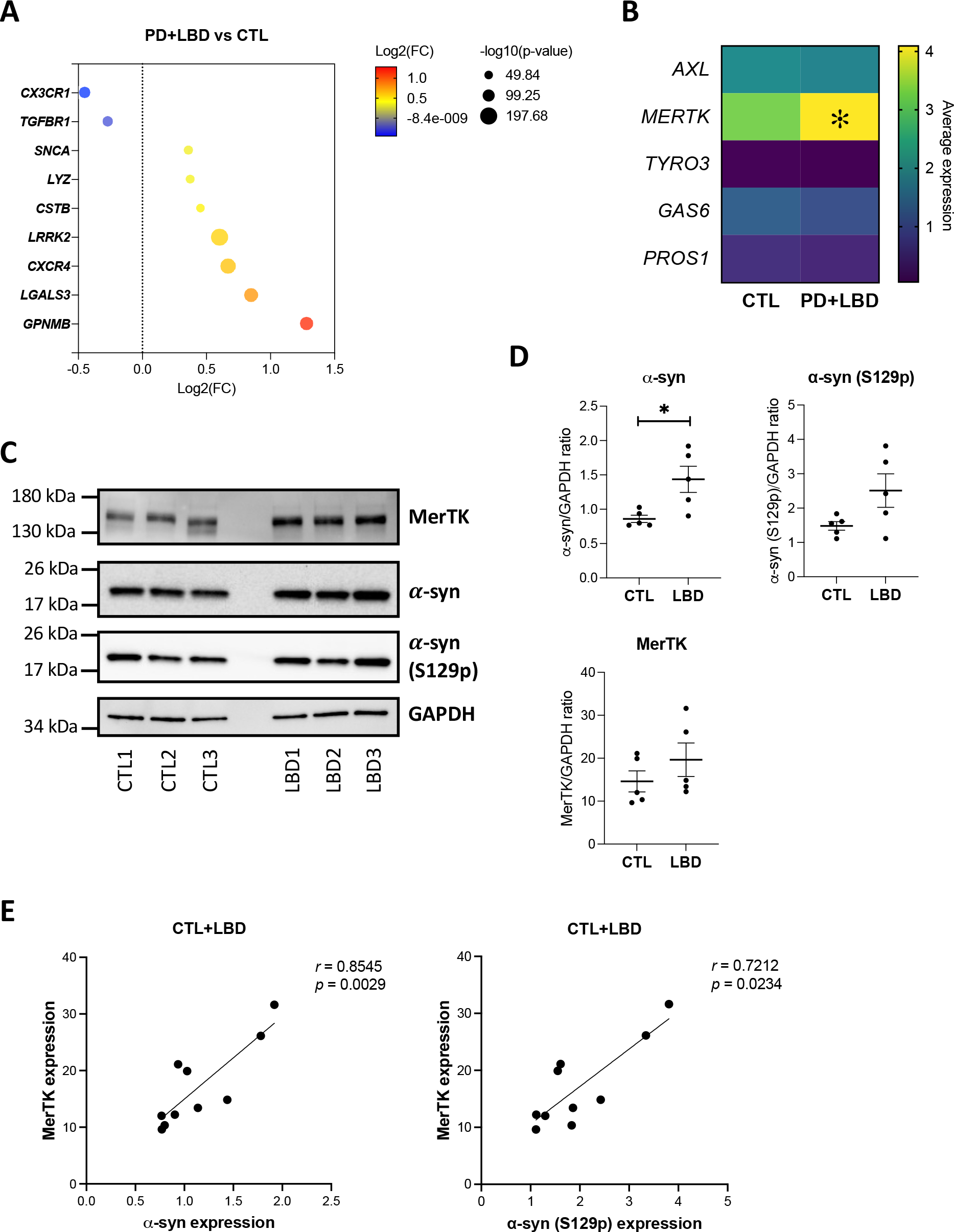
MerTK expression in Parkinson’s disease and Lewy body dementia. (A-B) snRNAseq data from Kamath *et al.*, 2022 were used to compare microglia from substantia nigra of seven Parkinson’s disease patients (PD) and three Lew body dementia patients (LBD) versus eight control donors (CTL). (A) Dotplot depicting examples of identified DEGs between PD/LBD and CTL. (B) Heatmap showing the average expression of genes encoding TAM receptors and their activating ligands. \**p* < 0.05 in DEG analysis. (C-E) Western blotting of MerTK, *α*-syn, phosphorylated *α*-syn (S129p) and GAPDH in human cortex of LBD and CTL donors. (C) Representative blots. Proteins were extracted in the presence of 5% SDS to solubilize *α*-syn aggregates. (D) Quantification of *α*-syn, phosphorylated *α*-syn (S129p) and MerTK expression. Mann-Whitney tests were performed. Mean +/- SEM of *n* = 5, \**p* < 0.05. (E) Correlation analysis of *α*-syn and MerTK expression across CTL and LBD samples (*n* = 10). ‘*r*’ denotes Spearman’s correlation coefficient.

Upregulation of MerTK protein expression in the context of synuclein accumulation was ascertained by Western blotting using cortex tissues from five patients with clinicopathological diagnosis of Lewy body dementia and five control donors (Supplementary Table 2). Significantly higher amounts of *α*-syn were detected in the cortex lysates of Lewy body dementia patients (Figure 5C-D). Although overall MerTK expression showed a non-significant increase in patient samples versus controls (Figure 5C-D), MerTK strongly correlated with *α*-syn levels (*p* = 0.0029; Figure 5E). MerTK expression also positively correlated with *α*-syn phosphorylated at serine 129 (S129p), an indicator of *α*-syn aggregation (*p* = 0.0234; Figure 5E). Altogether, data suggest an upregulation of MerTK expression in contexts of *α*-syn accumulation and aggregation.

## Discussion

In summary, our study identified MerTK as an essential receptor that mediates *α*-syn fibril uptake by human microglia. Using orthogonal assays, including pharmacological inhibition of MerTK kinase activity, *MERTK* downregulation induced by phenotypic polarization and *MERTK* knockdown by siRNA, we show that *α*-syn fibril internalization by human microglia is dependent on MerTK. MerTK-mediated internalization of *α*-syn PFFs was immunologically silent, failing to induce the expression and secretion of inflammatory cytokines. Moreover, *MERTK* mRNA expression was found to be increased in nigral microglia from Parkinson’s disease/Lewy body dementia patients compared to control donors, and MerTK protein expression in the human cortex positively correlated with *α*-syn level. Such findings support a role for MerTK-mediated phagocytosis in a setting of *α*-syn accumulation. Accordingly, burden analysis revealed a significant association between functionally deleterious, rare *MERTK* variants and Parkinson’s disease. This was true in the UKBB cohort of patients but not in the AMP-PD cohort, which could be a result of specific variants associated with Parkinson’s disease being underrepresented in the AMP-PD cohort. Replication of the burden analysis in future cohorts will be necessary to gain confidence that MerTK dysfunction is genetically associated with Parkinson’s disease.

Phagocytosis of cellular debris is a complex process that involves multiple cell surface proteins, some mediating tethering of the target and some initiating engulfment through cytoskeletal remodeling. Pattern recognition receptors such as TLRs can come into contact with their agonistic motifs during this process and trigger an inflammatory response.^54^ *α*- syn has been repeatedly observed to induce inflammatory cytokine release from murine microglia,^16–20^ albeit with some contradictory findings.^55^ In regards to human iMGL, Trudler *et al.*^56^ previously observed inflammasome activation and IL-1*β* secretion following cell treatment with non-sonicated large size *α*-syn PFFs (>2000 nm), whereas Rostami *et al.*^14^ failed to observe IL-1*β* secretion from iMGL following sonicated, smaller *α*-syn PFFs. In regards to hMGL, non-sonicated *α*-syn PFFs were previously shown by Pike *et al.*^57^ to induce inflammasome activation and subsequent IL-1*β* release from hMGL. In our study, *α*-syn PFFs sonicated to *∼*50-100 nm in size were employed because of the demonstrated pathogenicity and propensity for propagation of smaller PFFs.^58, 59^ These small PFFs were actively phagocytosed, but failed to induce IL-1*β* secretion from hMGL or iMGL. Notably, microglia in both Pike *et al.*’s. and Trudler *et al.*’s studies were cultured in the presence of granulocyte-macrophage colony-stimulating factor (GM-CSF), which might have influenced microglia response to *α*-syn PFFs. Finally, Pike *et al.* used saturating concentrations of *α*-syn PFFs (5-20 μM), which might have resulted in stress-induced non-specific activation of microglia.

Although the TAM receptors AXL and MerTK were thought to have redundant roles in phagocytosis, MerTK but not AXL has been demonstrated to mediate homeostatic phagocytosis. Daily clearance of photoreceptor outer segments by retinal pigment epithelial cells in the retina and apoptotic germ cells by Sertoli cells in the testis are mediated by MerTK,^60^ without which accumulation of cellular debris leads to tissue degeneration.^29, 61–63^ AXL and MerTK expression are also differentially regulated by tolerogenic and inflammatory stimuli: whereas MerTK expression is increased by treatment with glucocorticoids, AXL is upregulated and MerTK downmodulated following lipopolysaccharide or polyinosinic:polycytidylic treatment of macrophages.^64, 65^ Here we observed that microglial uptake of *α*-syn fibrils is dependent on MerTK but not AXL despite both receptors being expressed by human microglia. Consistent with the *in vitro* observations, only *MERTK* expression was found to be significantly higher in nigral microglia from Parkinson’s disease/Lewy body dementia patients compared to controls, and *MERTK* but not *AXL* variants were genetically associated with Parkinson’s disease cases in the UKBB cohort.

In a study that quantified the presence of *α*-syn inclusions in different cell types of the olfactory bulb of Parkinson’s disease patients, *∼*8% of microglia have been shown to contain inclusions, a strikingly high percentage considering that *∼*9% of neurons were found to contain inclusions in the same study.^66^ The main source of microglia-internalized *α*-syn in the Parkinson’s disease brain remains unclear. *α*-syn has been detected in the cerebrospinal fluid and brain interstitial fluid, suggesting the protein is released by neurons into the extracellular space through secretion or cell death.^67, 68^ The current study therefore focused on the role of MerTK in internalization of extracellular *α*-syn fibrils. However, MerTK could well be mediating *α*-syn internalization through phagocytosis of synapses^69^ and apoptotic neurons,^70^ limiting neuronal spread of *α*-syn.

MerTK has been receiving increasing attention as a potential therapeutic target in proteinopathies in the recent years. In the APP/PS1 mouse model of Alzheimer’s disease, Axl and MerTK have been shown to be important for microglia migration to amyloid plaques.^71^ Genetic ablation of *Axl* and/or *Mertk* led to accelerated death of the animals,^72^ indicative of a protective role for these receptors against disease progression. Longitudinal studies revealed that higher circulating levels of cleaved, soluble TAM receptors and their ligand GAS6 in the cerebrospinal fluid are associated with better cognitive outcome in Alzheimer’s disease.^73, 74^ To our knowledge, the current study represents the first *in vitro* demonstration of MerTK involvement in protein aggregate internalization. Pharmacological targeting of MerTK is difficult because of its wide expression and pleiotropic roles. Bispecific antibodies and hybrid proteins with affinities to both MerTK and amyloid beta have been designed to specifically target amyloid beta to MerTK-mediated, immunologically silent phagocytosis in Alzheimer’s disease.^75, 76^ Similar strategies could be adopted to enhance *α*-syn clearance in synucleinopathies.

In conclusion, we report for the first time a role for MerTK in the uptake of *α*-syn fibrils by human microglia. This pathway could be leveraged to promote *α*-syn fibril clearance in synucleinopathies such as Parkinson’s disease without inducing inflammatory response of microglia.

## Acknowledgements

We thank lab members Manon Blain, Qiao-Ling Cui, Genevieve Dorval, Andrea Krahn, Wolfgang Reintsch, Julien Sirois and Vincent Soubannier for technical or administrative assistance. We thank BDRL members Kimberly A. Aldinger, Dan Doherty, Ian G. Phelps, Jennifer C. Dempsey, Kevin J. Lee, and Lucinda A. Cort. We would like to thank the participants in the different cohorts for contributing to this study. This research used the NeuroHub infrastructure and was undertaken thanks in part to funding from the Canada First Research Excellence Fund, awarded through the Healthy Brains for Healthy Lives (HBHL) initiative at McGill University, Calcul Québec and Compute Canada. UKBB Resources were accessed under application number 45551. Data used in the preparation of this article were obtained from the AMP PD Knowledge Platform. For up-to-date information on the study, visit https://www.amp-pd.org. AMP PD – a public-private partnership – is managed by the FNIH and funded by Celgene, GSK, the Michael J. Fox Foundation for Parkinson’s Research (MJFF), the National Institute of Neurological Disorders and Stroke, Pfizer, Sanofi, and Verily. Genetic data used in preparation of this article were obtained from the Fox Investigation for New Discovery of Biomarkers (BioFIND), the Harvard Biomarker Study (HBS), the Parkinson’s Progression Markers Initiative (PPMI), the Parkinson’s Disease Biomarkers Program (PDBP), the International LBD Genomics Consortium (iLBDGC), and the STEADY-PD III Investigators. BioFIND is sponsored by MJFF with support from the National Institute for Neurological Disorders and Stroke (NINDS). The BioFIND Investigators have not participated in reviewing the data analysis or content of the manuscript. For up-to-date information on the study, visit michaeljfox.org/news/biofind. The HBS is a collaboration of HBS investigators (Co-Directors: Brigham and Women’s Hospital: Clemens R. Scherzer, Massachusetts General Hospital: Bradley T. Hyman; Investigators and Study Coordinators: Brigham and Women’s Hospital: Yuliya Kuras, Karbi Choudhury, Michael T. Hayes, Aleksandar Videnovic, Nutan Sharma, Vikram Khurana, Claudio Melo De Gusmao, Reisa Sperling; Massachusetts General Hospital: John H. Growdon, Michael A. Schwarzschild, Albert Y. Hung, Alice W. Flaherty, Deborah Blacker, Anne-Marie Wills, Steven E. Arnold, Ann L. Hunt, Nicte I. Mejia, Anand Viswanathan, Stephen N. Gomperts, Mark W. Albers, Maria Allora-Palli, David Hsu, Alexandra Kimball, Scott McGinnis, John Becker, Randy Buckner, Thomas Byrne, Maura Copeland, Bradford Dickerson, Matthew Frosch, Theresa Gomez-Isla, Steven Greenberg, Julius Hedden, Elizabeth Hedley-Whyte, Keith Johnson, Raymond Kelleher, Aaron Koenig, Maria Marquis-Sayagues, Gad Marshall, Sergi Martinez-Ramirez, Donald McLaren, Olivia Okereke, Elena Ratti, Christopher William, Koene Van Dij, Shuko Takeda, Anat Stemmer-Rachaminov, Jessica Kloppenburg, Catherine Munro, Rachel Schmid, Sarah Wigman, Sara Wlodarcsyk; Data Coordination: Brigham and Women’s Hospital: Thomas Yi; Biobank Management Staff: Brigham and Women’s Hospital: Idil Tuncali.) and funded through philanthropy and National Institutes of Health (NIH) and Non-NIH funding sources. The HBS Investigators have not participated in reviewing the data analysis or content of the manuscript. PPMI – a public-private partnership – is funded by MJFF and funding partners, including 4D Pharma, AbbVie Inc., AcureX Therapeutics, Allergan, Amathus Therapeutics, Aligning Science Across Parkinson’s (ASAP), Avid Radiopharmaceuticals, Bial Biotech, Biogen, BioLegend, Bristol Myers Squibb, Calico Life Sciences LLC, Celgene Corporation, DaCapo Brainscience, Denali Therapeutics, The Edmond J. Safra Foundation, Eli Lilly and Company, GE Healthcare, GSK, Golub Capital, Handl Therapeutics, Insitro, Janssen Pharmaceuticals, Lundbeck, Merck & Co., Inc., Meso Scale Diagnostics, LLC, MJFF, Neurocrine Biosciences, Pfizer Inc., Piramal Imaging, Prevail Therapeutics, F. Hoffmann-La Roche Ltd and its affiliated company Genentech Inc., Sanofi Genzyme, Servier, Takeda Pharmaceutical Company, Teva Neuroscience, Inc., UCB, Vanqua Bio, Verily Life Sciences, Voyager Therapeutics, Inc., Yumanity Therapeutics, Inc.. The PPMI Investigators have not participated in reviewing the data analysis or content of the manuscript. For up-to-date information on the study, visit www.ppmi-info.org. PDBP consortium is supported by the NINDS at the National Institutes of Health. A full list of PDBP investigators can be found at https://pdbp.ninds.nih.gov/policy. The PDBP investigators have not participated in reviewing the data analysis or content of the manuscript. Genome Sequencing in Lewy Body Dementia and Neurologically Healthy Controls: A Resource for the Research Community.” was generated by the iLBDGC, under the co-directorship by Dr. Bryan J. Traynor and Dr. Sonja W. Scholz from the Intramural Research Program of the U.S. National Institutes of Health. The iLBDGC Investigators have not participated in reviewing the data analysis or content of the manuscript. For a complete list of contributors, please see: bioRxiv 2020.07.06.185066; doi: https://doi.org/10.1101/2020.07.06.185066. STEADY-PD III is a 36-month, Phase 3, parallel group, placebo-controlled study of the efficacy of isradipine 10 mg daily in 336 participants with early Parkinson’s Disease that was funded by the NINDS and supported by MJFF and the Parkinson’s Study Group. The STEADY-PD III Investigators have not participated in reviewing the data analysis or content of the manuscript. The full list of STEADY PD III investigators can be found at: https://clinicaltrials.gov/ct2/show/NCT02168842.

## Supplementary material

Supplementary material is available at *Brain* online.

