## Supplementary Fig. for "MerTK mediates the immunologically silent uptake of alpha-synuclein fibrils by human microglia"

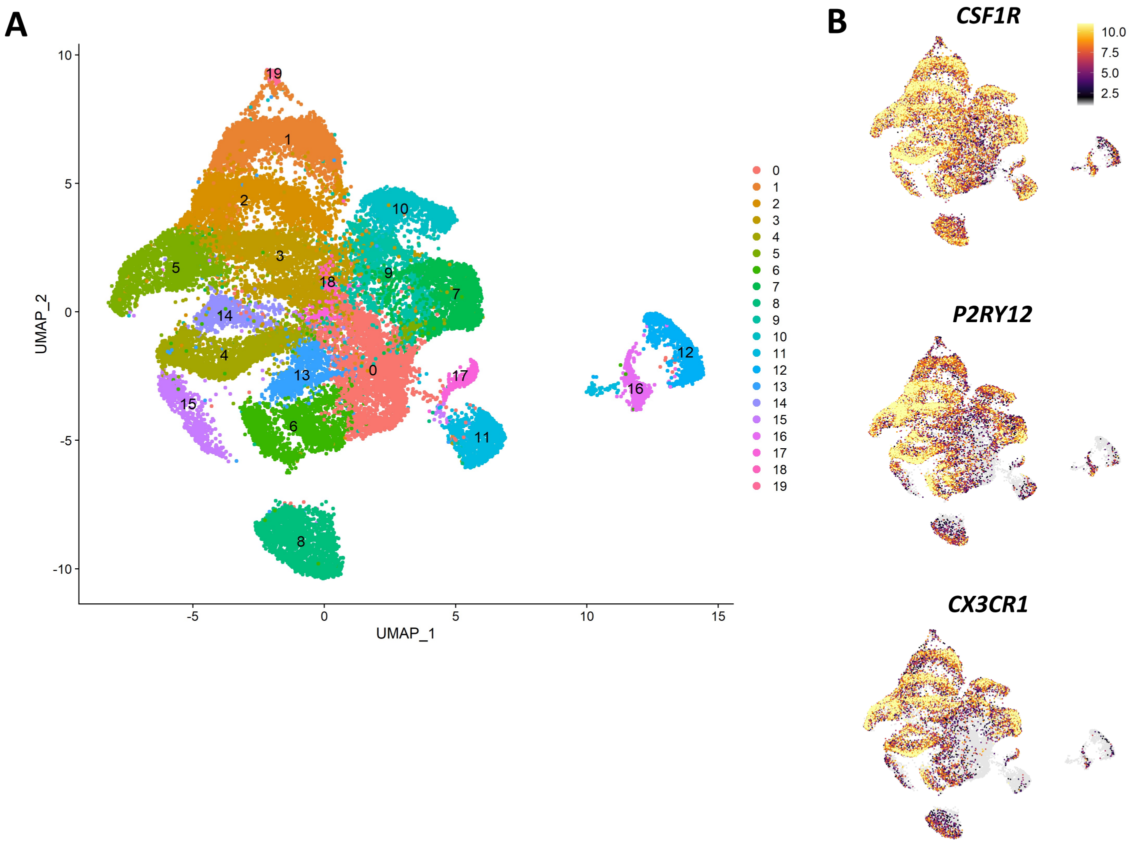


**Supplementary Figure 1. Microglia population identified by snRNAseq.** (A) uniform manifold approximation and projection (UMAP) of microglia clusters identified in the snRNAseq data from Kamath *et al.*, 2022. (B) Expression of microglia canonical markers in the identified clusters. Color scale represents gene expression level.

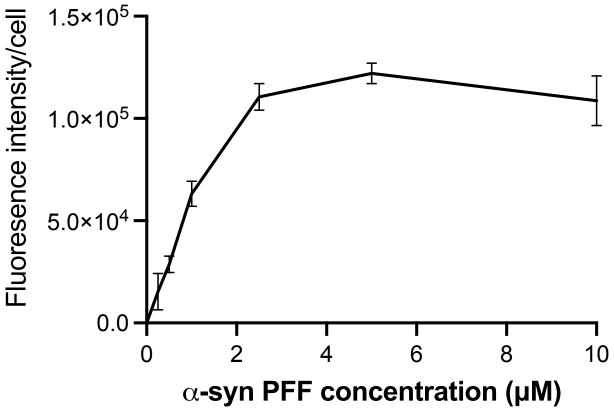


**Supplementary Figure 2. Green fluorescence intensity in function of pHrodo™ Green-labelled α-syn PFF concentration.** Quantification of green fluorescence intensity per cell in iMGL challenged with pHrodo™ Green-labelled α-syn PFFs for two hours. Mean +/- SEM of three technical replicates are presented.

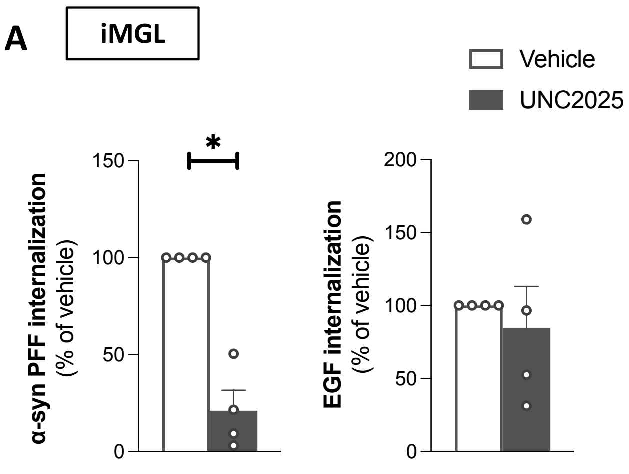


**Supplementary Figure 3. Effect of UNC2025 on microglial uptake of Alexa Fluor 488-labelled α-syn PFFs.** iMGL were pretreated with vehicle or UNC2025 (3 μM) for one hour and then challenged with Alexa Fluor 488-labelled α-syn PFFs or EGF for two hours. (A) Quantification of green fluorescence intensity per cell. Data were normalized to the vehicle-treated conditions. A Mann-Whitney test was performed. Mean +/- SEM of $n = 4$, *$p < 0.05$. (B) Representative fluorescence images of iMGL counterstained with Hoechst 33342 (blue). Scale bar = 150 $\mu m$.

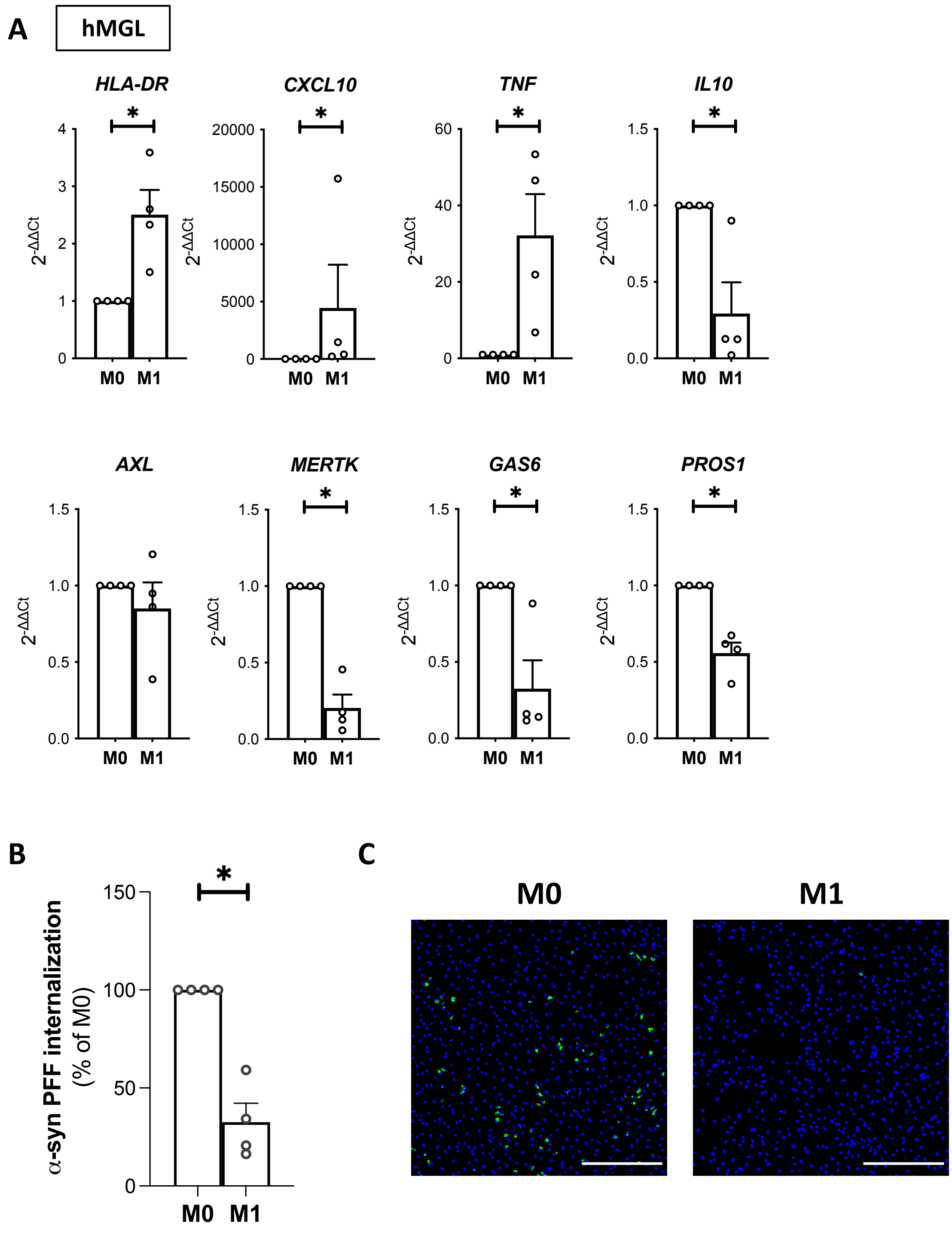


**Supplementary Figure 4. Effect of M1 polarization on hMGL ability to internalize α-syn PFFs.** hMGL were M1-polarized using IFNγ (20 ng/mL) and $Pam_3CSK_4$ (100 ng/mL) treatment for 48 hours or left unpolarized (M0 hMGL). (A) qRT-PCR data. Mann-Whitney tests were performed. Mean +/- SEM of $n$ = 4, $*p < 0.05$. (B) Quantification of green fluorescence intensity per cell in hMGL culture challenged with pHrodo™ Green-labelled α-syn PFFs for two hours. Data were normalized to the M0 conditions. A Mann-Whitney test was performed. Mean +/- SEM of $n = 4$, $*p < 0.05$. (C) Representative fluorescence images of cells challenged with pHrodo™ Green-labelled α-syn PFFs for two hours and counterstained with Hoechst 33342 (blue). Scale bar = 300 $\mu m$.

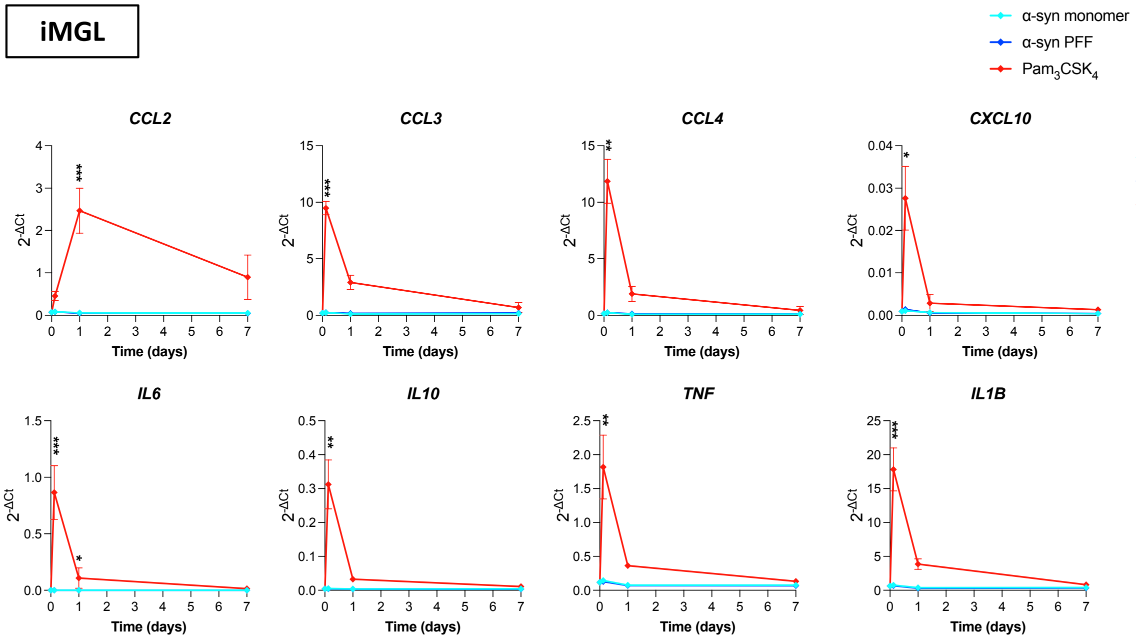


**Supplementary Figure 5. Effect of α-syn PFF treatment on cytokine expression.** iMGL were treated with α-syn monomers (1 μM), α-syn PFFs (1 μM) or $Pam_3CSK_4$ (100 ng/mL) for three hours, one day or seven days. Expression of cytokine genes were assessed by qRT-PCR. Kruskal-Wallis tests were performed, followed by Dunn's post hoc test. Mean +/- SEM of $n = 5$, *$p < 0.05$, **$p < 0.01$, ***$p < 0.001$.
